# Dysregulation of the CeA→BNST pathway is associated with alcohol-induced anxiety-like behaviors in female mice

**DOI:** 10.64898/2026.11.20.768212

**Authors:** Sarah Chong, Jobe Ritchie, Lisa Taxier, Sloan Markowitz, Lili Kooyman, Maya Eberle-Redding, Chloe Gallagher, Madison Campeau, Thomas Kash, Alison Roland

## Abstract

The central amygdala (CeA) and bed nucleus of stria terminalis (BNST) constitute the major output regions of the extended amygdala, a key neural substate of anxiety that drives heightened negative affect during alcohol withdrawal. In keeping with clinical evidence demonstrating that stronger amygdala-BNST functional connectivity positively correlates with higher anxiety measures in individuals with Alcohol Use Disorder (AUD), a growing body of preclinical evidence implicates a role for the CeA BNST pathway in mediating alcohol withdrawal-induced anxiety. However, little is known about how the CeA and BNST interact during alcohol abstinence and anxiety. The dysregulation of kappa opioid receptor (KOR) signaling in the amygdala is strongly implicated in AUD, and KOR function in the CeA and BNST is linked to anxiety and alcohol dependence. Yet, whether alcohol dependence alters CeA BNST connectivity, and how KOR signaling contributes to these circuit adaptations remains unknown. Using fiber photometry, we found that chronic intermittent ethanol (CIE) – a translational model of alcohol dependence – increased anxiety-like behaviors in female mice and BNST activity during anxiogenic contexts. Moreover, CIE-exposed females exhibited a frequency-specific increase in CeA-BNST coherence during an anxiogenic task. Our circuit-specific electrophysiological studies identified a female-specific postsynaptic disruption in the CeA→BNST circuit following alcohol dependence, as well as the first evidence that KORs suppress CeA→BNST connectivity in female mice. Together, these findings demonstrate that CIE promotes BNST hyperactivity and frequency-specific CeA-BNST coherence during anxiogenic behaviors in female mice and identify discrete, sex-specific adaptations in the CeA BNST circuit that may contribute to BNST hyperactivity during alcohol withdrawal.

## Introduction

Alcohol use disorder (AUD) is a leading cause of preventable death worldwide.^1^ Anxiety disorders are highly prevalent in individuals with AUD,^2,3^ and their comorbidity worsens symptoms of either condition, limiting treatment and recovery outcomes.^4,5^ While there are three FDA-approved medications for AUD, no single pharmacotherapeutic has been successful in treating all AUD patients, and none treats negative affect specifically.^5–7^ Importantly, the prevalence of AUD in women has risen sharply in recentdecades,^8,9^ and clinical evidence demonstrates that women are more likely than men to experience anxiety and to consume alcohol to alleviate negative affect.^10,11^ Although heightened negative affect during alcohol dependence and withdrawal is a major contributor to relapse in AUD, the neural circuit mechanisms driving these affective changes are poorly understood.^2,12^ Elucidating such mechanisms will inform the development of more efficacious therapeutics for AUD.

The extended amygdala, including the central amygdala (CeA) and bed nucleus of stria terminalis (BNST), is a key neural substrate of anxiety-related behavior and avoidance.^12–15^ Both regions contain diverse populations of various neuronal subtypes, most of which express the inhibitory neurotransmitter GABA.^16^ Neuronal activity is increased in both the CeA and BNST during alcohol withdrawal,^17–19^ and heightened activity in these regions drives anxiety-like behaviors.^13,20–24^ Consistent with a role for this circuitry in AUD, clinical studies demonstrate increased amygdala-BNST functional connectivity that positively correlates with anxiety measures in individuals with AUD.^11,15^ Given that the CeA strongly innervates the BNST,^25,26^ this suggests that the CeA BNST pathway plays a key role in mediating alcohol withdrawal-induced anxiety-like states. The dysregulation of distinct peptide neuromodulator systems (e.g., dynorphin and kappa opioid receptors) has also been strongly implicated in the etiology of AUD and mood disorders characteristic of heightened avoidance.^12,27–29^ Consistent with this, kappa opioid receptor (KOR) function in the CeA and BNST is linked to anxiety-like behavior.^29–32^ However, whether alcohol dependence alters CeA BNST connectivity, and how KOR signaling contributes to these circuit adaptations, remains unknown.

To address the knowledge gap in mechanisms underlying alcohol-induced negative affect, the present study evaluated *in vivo* neuronal dynamics in the CeA and BNST of mice during tests of anxiety-like behavior following chronic intermittent ethanol, a preclinical model of alcohol dependence. We next used optogenetics and slice electrophysiology to assess whether alcohol dependence alters synaptic connectivity and KOR modulation in the CeA BNST pathway.

## Methods and Materials

### Animals

Experiments were performed on adult male and female mice 8-17 weeks of age at the beginning of the experiments. C57BL/6J mice (Stock #000664, Jackson Laboratories, Bar Harbor, ME) arrived at the vivarium at 7-8 weeks of age and acclimated for at least 1 week prior to stereotaxic surgery. VGAT-ires-Cre mice were generated as described previously.^33^ Animals were group housed in a ventilated and temperature-controlled vivarium on a standard 12:12 hour light-dark cycle with *ad libitum* access to rodent chow and water. All animal procedures were performed in accordance with the NIH guidelines for animal research and with the approval of Institutional Animal Care and Use Committee at the University of North Carolina at Chapel Hill.

### Stereotaxic Surgeries

Mice were placed in a stereotaxic frame (Kopf Instruments, Tujunga, CA) and adeno-associated virus (AAV7-hsyn-GCaMP8m-EYFP-WPRE or AAV5-Ef1α-DIO-ChR2(H134R)-EYFP) was infused into either the CeA (M/L ±2.90, A/P -1.20, D/V -4.70) or BNST (M/L ±0.90, A/P 0.30, DV -4.35) at a flow rate of 100 nL/min and left in place for at least 5 minutes to ensure viral diffusion. Fiber optic cannulae (200 uM, 0.37 nA, Neurophotometrics, San Diego, CA) were then implanted in the contralateral CeA and BNST using the same coordinates as for viral injections and secured using C&B-Metabond (Parkell, Inc., Edgewood, NY), with mice counterbalanced for side placement.

### Chronic Intermittent Ethanol (CIE)

Chronic intermittent ethanol exposure was achieved via vapor inhalation as previously described.^34,35^

### Fiber Photometry

Fiber photometry analysis was conducted using open-source software (Tracy; https://github.com/JobeRitchie/TRACY-Photometry-Processing-Suite). Photometry was acquired with alternating 470 nm and 415 nm channels at a sampling rate of 30 Hz per channel. Traces were deinterleaved and corrected for photobleaching by fitting a biexponential function. Motion artifacts were removed by regressing the 415 nm ΔF/F onto the 470 nm ΔF/F using robust Huber regression and subtracting the fitted isosbestic component. Corrected traces were z-scored across session then lowpass filtered with zero-phase second-order Butterworth filter (6 Hz cutoff). Photometry was aligned to behavior using the nearest-match computer time stamp at start of recording. Spectral coherence between CeA and BNST was conducted as previously described.^36,37^

### Behavioral Analysis

Behavior analysis for the EPM and NSF assays was conducted using ABEL v0.14.0 (Active-learning Behavior Estimation and Labeling), an open-source pipeline for supervised machine learning (https://github.com/JobeRitchie/ABEL).

### Elevated Plus Maze

Elevated Plus Maze (EPM) testing was conducted 24 hours after the last cycle of CIE as previously described.^38^ Mice were allowed to freely explore during a 10-minute session. Only female mice were run in this experiment due to a logistical oversight.

### Novelty Suppressed Feeding Test

In the week prior to behavioral testing, mice were provided with access to a single pellet of high-fat chow (60% fat, Research Diets D12492, New Brunswick, NJ) in their home cage to familiarize them to the food prior to testing. Mice were subjected to mild food restriction for 2-4 hours immediately prior to the test.

During the test session mice were placed into a brightly lit arena (51×51 cm) that contained a single high-fat pellet in the center. Mice were allowed to explore the arena freely for ten minutes.

### Whole-Cell Voltage-Clamp Recordings

5-7 days after the last cycle of CIE, mice were sacrificed for slice electrophysiology and recordings performed as previously described.^33^ For kinetic analysis of sIPSCs, the first 30 monosynaptic events with the following criteria were selected from each recording: single event, stable baseline, sharp rise, and exponential decay.^39^ No more than 2 cells per animal were included for analysis. U69593 was purchased from Sigma-Aldrich (St. Louis, MO) and dissolved in 0.1 M HCl. All other drug information is located in **Supplement 1**.

### Statistical Analysis

Data are expressed as means ± SEM. Statistical analyses were performed using GraphPad Prism 10.6.1 Software (La Jolla, CA). Differences between groups were analyzed using t-tests, 1-way ANOVA, 2-way ANOVA, and 3-way ANOVA as appropriate. Sidak’s post-hoc test was used following significance with an ANOVA. Correlations were evaluated using Pearson’s correlation coefficient. For NSF, Kaplan-Meier survival analysis was used to analyze feeding latencies due to the non-normal distribution of the data and the presence of censored observations from mice that did not eat during the 10-minute period. The Mantel-Cox logrank test was used to assess differences between treatment groups. Significance was set at p < 0.05. Changes in synaptic transmission were assessed with either a 2-way ANOVA or a two-sample *t* test with Welch’s correction where appropriate. Effects of U69593 during electrophysiological recordings were evaluated by comparing the magnitude of peak oIPSC amplitude between the baseline and washout (when drug had reached maximal effect) periods using *t* tests. Mice with off-target virus expression or fiber placements were excluded from analysis.

## Results

### Females exhibit elevated BNST calcium signal and CeA-BNST coherence during anxiety-like behaviors after CIE

We first sought to understand how chronic intermittent ethanol (CIE) exposure, a model of alcohol dependence,^34,35^ impacts neuronal activity in the CeA and BNST during the elevated plus maze test (EPM), a pharmacologically validated assay used for measuring anxiety-related behaviors. Fiber photometry recordings of CeA and BNST calcium dynamics were acquired while female mice freely explored the EPM, beginning 24 hours following the last CIE exposure **(Fig. 1A-C)**, a timepoint at which CIE was previously shown to promote heightened anxiety-like behavior.^40^ We used a machine learning approach to characterize discrete, ethologically relevant behaviors during the assay, including protected head-dips, stretch-attend postures, freezing, rearing, and grooming **(Fig. 1B)**. Mean blood alcohol concentrations per CIE cycle were maintained near and above binge intoxication levels **(Fig. 1D).**

**Figure 1.**
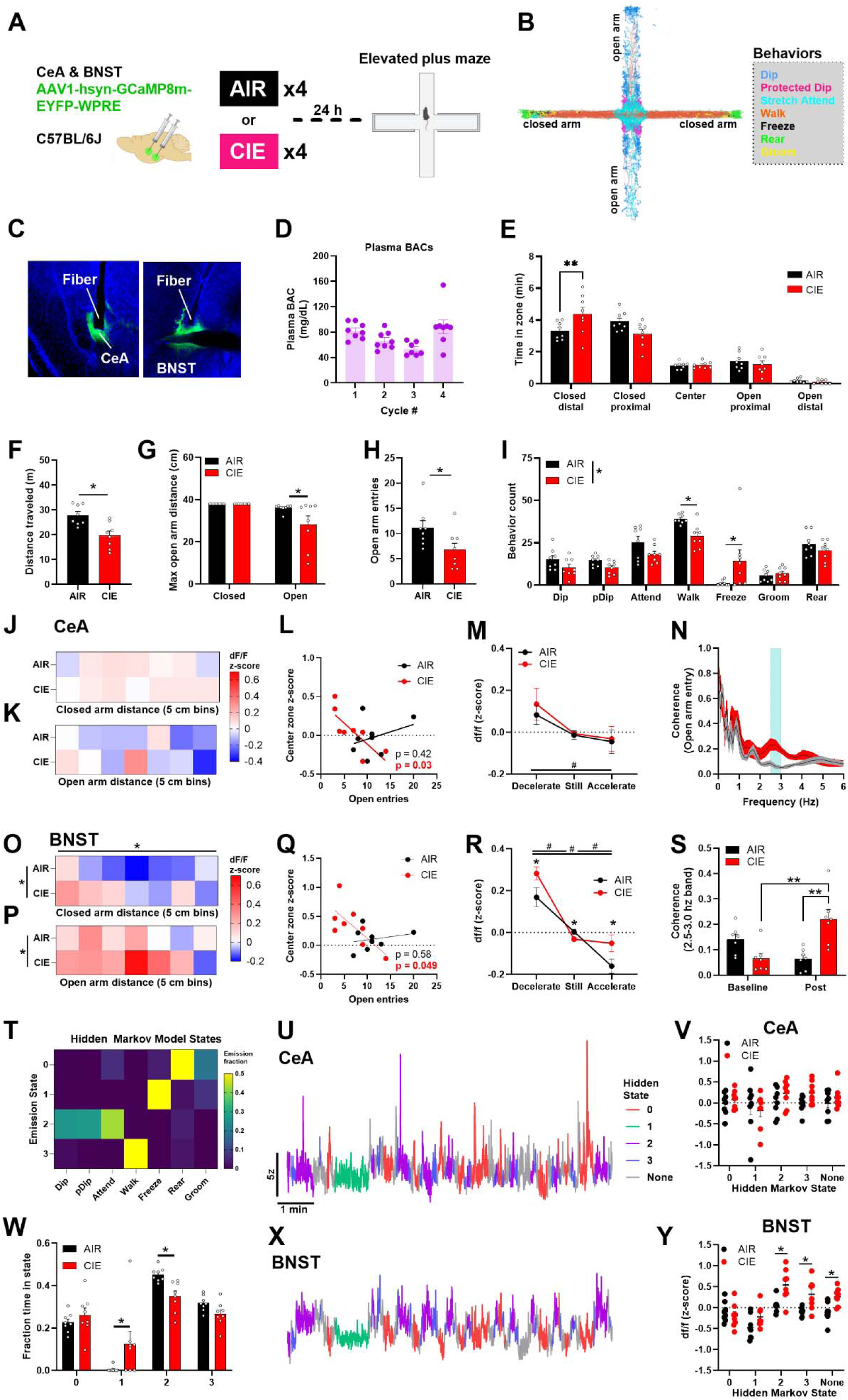
Females exhibit anxiety-like behavior and elevated BNST calcium signal during surveillance behaviors after CIE. **A)** Schematic of timeline for EPM fiber photometry experiments. Female C57BL/6J mice received GCaMP and fiber optic implantations into the CeA and BNST, in parallel per mouse, prior to CIE exposure. Mice underwent exposure to either vaporized ethanol (CIE; N = 8 female) or air (AIR; N = 8 female) on a 16hr-ON/8hr-OFF cycle for four weeks. Fiber photometry recordings of GCaMP were acquired while mice explored the EPM, beginning 24 hours after the last vapor exposure. **B)** Heat-map depicting spatial locations of ethological behaviors characterized by ABEL in the EPM apparatus. **C)** Histological verification of virus and optic fiber placement in CeA and BNST. **D)** Mean plasma blood alcohol concentrations per cycle were maintained near and above binge intoxication levels (mean BAC = 72.8 ± 7.88 mg/dL; Cycle 1: 82.2 ± 4.51 mg/dL; Cycle 2: 66.8 ± 4.81 mg/dL; Cycle 3: 53.6 mg/dL ± 3.90 mg/dL; Cycle 4: 88.6 mg/dL ± 10.9 mg/dL). **E)** CIE increased time spent in the distal portions of the closed arms; 2-way repeated measures ANOVA, location x treatment interaction effect, F_(4,56)_ = 3.911, p = 0.007; AIR closed distal < CIE closed distal, p = 0.004. **F)** CIE reduced distance traveled (two-tail t-test, t_(13.33)_ = 3.56, p = 0.003). CIE reduced the maximal distance travelled into the open-arms **(G;** 2-way ANOVA, arm x treatment interaction effect, F_(1,28)_ = 3.78, p = 0.06; AIR max distance in open arm > CIE max distance, p = 0.02), as well as entries made into the open-arms **(H;** two-tailed t-test, t_(13.95)_ = 2.23, p = 0.04). **I)** CIE increased freezing bouts and reduced walking bouts (2-way ANOVA, behavior x treatment interaction effect, F_(6,98)_ = 4.39, p = 0.001; AIR freeze < CIE freeze, p = 0.004; AIR walk > CIE walk, p = 0.04), with a trend for overall reduction of all other assayed behavioral bouts (2-way ANOVA, treatment main effect, F_(1,98)_ = 2.91, p = 0.09). **J-M:** Data obtained from CeA. CeA calcium signal had no significant difference across the spatial extent in the closed **(J)** and open **(K)** arms, independent of CIE exposure. **L)** CeA calcium activity in the center zone negatively correlated with number of open-arm entries in CIE mice (R^2^ = 0.60, p = 0.03), with no significant relationship in AIR mice (R^2^ = 0.11, p = 0.42). **M)** CeA calcium activity decreased during acceleration relative to deceleration, independent of CIE exposure; acceleration main effect, F_(2,42)_ = 4.82, p = 0.01; acceleration < deceleration, p = 0.02. **O-R:** Data acquired from BNST. BNST calcium signal was increased in the distal-most portion of the closed arm **(O)** relative to the proximal extent (distance main effect, F_(6, 98)_ = 3.710, p < 0.05), and CIE increased calcium signal in closed **(O;** 2-way ANOVA, treatment main effect, F_(1,98)_ = 12.21, p < 0.001) and open arms **(P;** treatment main effect, F_(1,87)_ = 4.55, p = 0.04). **Q)** BNST calcium activity in the center zone negatively correlated with number of open-arm entries in CIE mice (R^2^ = 0.50, p < 0.05), with no significant correlation in AIR mice (R^2^ = 0.06, p = 0.57). **R)** BNST calcium activity decreased during acceleration relative to deceleration (acceleration main effect, F_(2,42)_ = 58.11, p < 0.001, acceleration < still < decleration, all p < 0.05) and was elevated in CIE mice; treatment x acceleration interaction effect, F_(2,42)_ = 3.59, p = 0.04; AIR acceleration < CIE acceleration, p < 0.02, AIR deceleration < CIE deceleration, p < 0.02. CIE increased CeA-BNST coherence in the 2.5-3.0 Hz frequency band **(N, S)** during open-arm entry bouts; 2-way ANOVA, time x treatment interaction effect, F_(1,24)_ = 20.61, p < 0.001; AIR post bout < CIE post bout, p = 0.001, CIE baseline < CIE post bout, p = 0.001. **T)** Heat-map depicting behavioral states comprised of ethological behaviors emitted by HMM analysis. Behavoral state mapped on to calcium signals in both the CeA **(U)** and BNST **(X)**. **V)** CIE had no effect on HMM state-associated signal in the CeA; 2-way ANOVA, all p > 0.05. **W)** CIE increased time spent in state 1 and reduced time spent in state 2; 2-way ANOVA, treatment x state interaction, F_(3,56)_ = 6.03, p = 0.001; AIR state 1 < CIE state 1, p = 0.02, AIR state 2 > CIE state 2, p = 0.05. **Y)** CIE increased BNST signal during states 2 and 3, as well as when not in a state; treatment x state interaction effect, F(3,56) = 6.03, p = 0.001; AIR state 2, 3, none < CIE state 2, 3, none, all p < 0.05.

CIE-exposed females exhibited behavior consistent with a hypolocomotive, anxiety-like state relative to alcohol-naïve females. CIE reduced the total distance traveled **(Fig. 1F)**, and analysis of discrete behaviors revealed a trend for a reduction in all behaviors assessed and a significant reduction in ambulation after CIE **(Fig. 1I)**. Importantly, we also found that CIE increased anxiety-like behaviors, as indicated by increased time in the distal portions of the closed arms **(Fig. 1E)**, decreased maximum distance travelled into the open arms **(Fig. 1G)** and reduced number of entries into the open arms **(Fig. 1H)**. Likewise, analysis of discrete behaviors revealed a significant increase in freezing bouts following CIE **(Fig. 1I)** consistent with a heightened anxiety-like state.

To assess whether these behavioral changes coincide with neuronal activity changes following CIE, we next examined calcium signal in the CeA and BNST across the spatial extent of the EPM. In the CeA, we found no effect of distance across the closed arms **(Fig. 1J)** or open arms **(Fig. 1K)** on calcium signal in either alcohol-naïve or CIE conditions. However, CIE mice exhibited a significant negative correlation between CeA activity in the center zone and the number of entries made into the open arms **(Fig. 1L)**, a correlation absent in alcohol-naïve mice **(Fig. 1L)**, suggesting that CIE-induced alterations in CeA neuronal activity promote behavioral avoidance. In the BNST, calcium signal scaled with proximity to the center zone during closed-arm locomotion, with lower signal in the distal-most part of the closed arms relative to the proximal part **(Fig. 1O)** independent of CIE exposure, demonstrating that BNST activity scales with the anxiogenic context. CIE exposure increased BNST calcium activity during locomotion in both the closed arms **(Fig. 1O)** and open arms **(Fig. 1P)**, demonstrating that CIE promotes BNST hyperactivity coincident with heightened anxiety-like behavior in mice. Similarly, CIE mice exhibited a significant negative correlation between BNST calcium activity in the center-zone and the number of open-arm entries **(Fig. 1Q)**, which was absent in AIR mice **(Fig. 1Q)**, lending further support to our hypothesis that CIE-induced increases in BNST activity promote heightened anxiety-like states in female mice.

To assess how calcium dynamics in CeA and BNST map on to the animals’ behavioral state, we investigated underlying behavioral states using hidden Markov modeling (HMM) analysis. Our analysis discovered 4 distinct behavioral states **(Fig. 1T)** comprised of the following behaviors: rearing and grooming (state 0), freezing (state 1), surveillance behaviors (dips, protected dips, stretch-attend postures; state 2), and ambulation (state 3). We found that CIE mice spent significantly more time in State 1 and less time in State 2 compared to AIR mice **(Fig. 1W),** consistent with a hypolocomotive state. All four of these distinct behavioral states mapped on to calcium signal in both the CeA **(Fig. 1U)** and BNST **(Fig. 1X)**. In the CeA, we found that CIE had no effect on state-specific calcium signals **(Fig. 1V)**. In the BNST, however, CIE mice exhibited elevated calcium signal during states 2 and 3 and when not in a specific state, relative to AIR mice **(Fig. 1Y)**, suggesting that BNST hyperactivation may promote CIE-induced behavioral changes.

We also assessed calcium activity specifically during acceleration and deceleration, as well as during a static locomotive state characterized by an absence of change in speed. We observed that CeA calcium activity was decreased during acceleration relative to deceleration **(Fig. 1M)**, independent of CIE exposure. In the BNST, calcium activity was similarly reduced during acceleration relative to deceleration and static states **(Fig. 1R)**, independent of CIE exposure. CIE exposure increased BNST calcium activity during acceleration and attenuated the decrease in activity during deceleration **(Fig. 1R)**.

Building on a growing body of evidence that implicates a role for increased CeA-BNST functional connectivity in alcohol-induced anxiety,^13,15,18,23,26,36,41^ we next examined CeA-BNST spectral coherence during anxiety-like behaviors. We assessed CeA-BNST spectral coherence during open arm entry bouts with a minimum open arm time of 10 seconds **(Fig. 1N)** and detected a significant increase in coherence in the 2.5-3.0 Hz frequency band in CIE mice relative to AIR mice **(Fig. 1S)**. Together, these findings point to a role for increased frequency-specific coherence between the CeA and BNST in promoting anxiety-related behaviors following CIE in female mice.

### Increased CeA and BNST activation during approach following CIE is associated with sex-specific differences in latency to feed in a novel context

Based on the findings in EPM, we examined the impact of CIE on behavior and signaling during the novelty-suppressed feeding test (NSF) at a later timepoint. We used fiber photometry to record real-time neuronal activity in the CeA and BNST of female and male mice during the NSF on day 5 of alcohol withdrawal **(Fig. 2A)**, a timepoint at which CIE was previously shown to promote heightened anxiety-like behavior.^40^ Mean plasma blood alcohol concentrations per cycle were maintained above binge intoxication levels **(Fig. 2B).**

**Figure 2.**
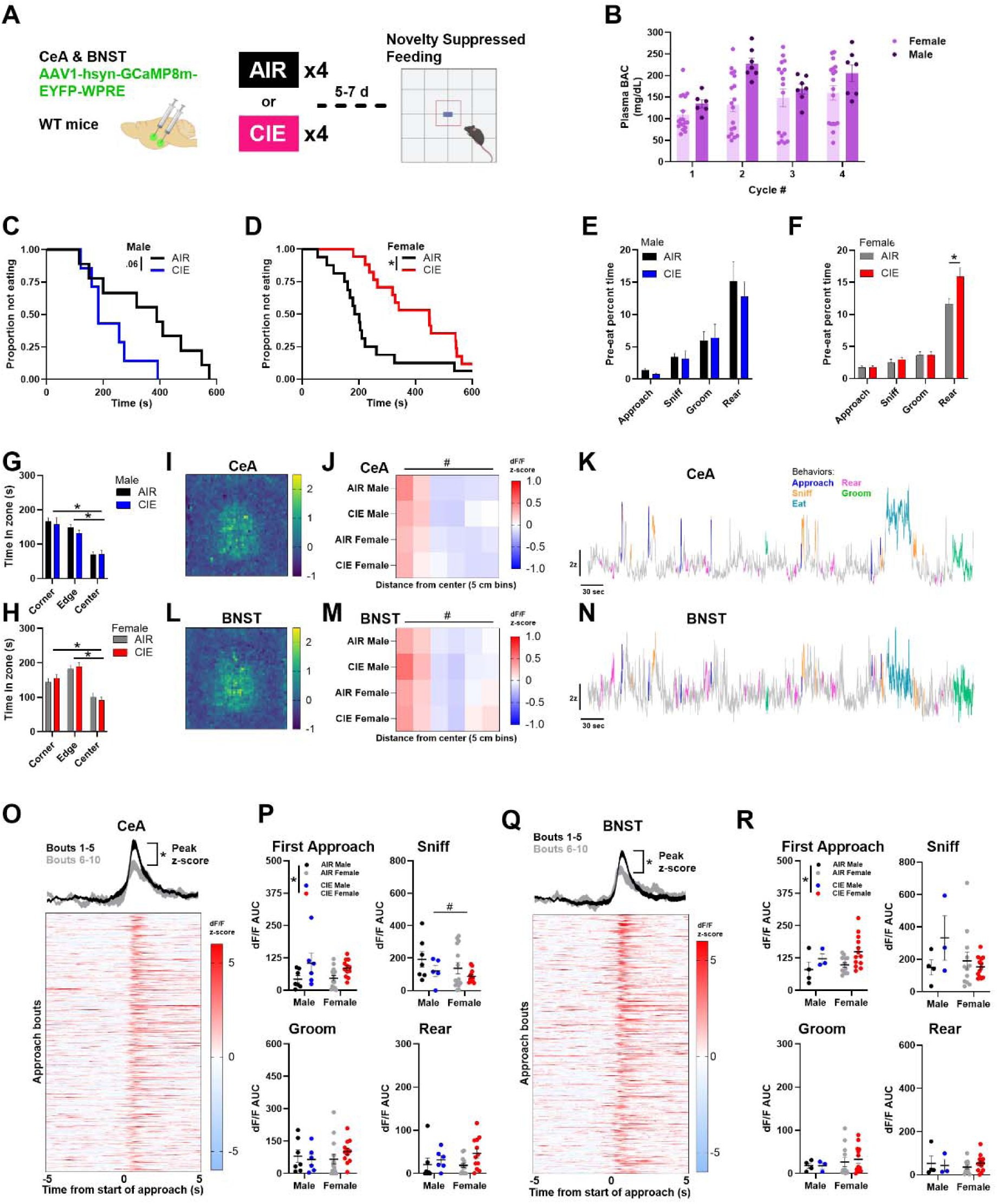
Increased CeA and BNST activation following CIE has sex specific effects on latency to feed in a novel context. **A)** Schematic of timeline for NSF fiber photometry experiments. Female and male C57BL/6J mice received GCaMP and fiber optic implantations into the CeA and BNST, in parallel per mouse, prior to CIE exposure. Mice underwent exposure to either vaporized ethanol (CIE; N = 16 female, 7 male) or air (AIR; N= 15 female, 8 male) on a 16hr-ON/8hr-OFF cycle for four weeks. Fiber photometry recordings of GCaMP were acquired while mice explored the NSF apparatus, beginning 5 days after the last vapor exposure. **B)** Mean plasma blood alcohol concentrations per cycle were maintained above binge intoxicaton levels in both females (mean = 137 ± 11.0 mg/dL; Cycle 1, mean = 109 ± 9.49 mg/dL; Cycle 2, mean = 132 ± 16.1 mg/dL; Cycle 3, mean = 148 ± 20.7 mg/dL; Cycle 4, mean = 160 ± 16.8 mg/dL) and males (mean = 184 ± 20.4 mg/dL; Cycle 1, mean = 135 ± 9.61 mg/dL; Cycle 2, mean = 228 ± 13.1 mg/dL; Cycle 3, mean = 169 ± 11.6 mg/dL; Cycle 4, mean = 206 ± 20.0 mg/dL). **C)** CIE males trended towards a reduction in latency to feed; Mantel-Cox test on survival curve, χ² = 3.67, p = 0.055. **D)** CIE increased latency to feed in females; χ² = 7.96, p = 0.005. **E)** Males exhibited no effect of CIE on time spent engaged in food-approach, sniffing, grooming, and rearing behaviors prior to eating; two-tailed t-tests, p > 0.05. **F)** Females exhibited increased time spent rearing prior to eating following CIE (t_(24.21)_ = 2.69, p = 0.01), with no effects of CIE on food-approach, sniffing, and grooming behaviors (two-tailed t-tests, p > 0.05). Both male **(G)** and female **(H)** mice spent less time in the anxiogenic center relative to periphery of the arena, and CIE had no effect on time spent in the center; 2-way ANOVA, main effect of zone, females: F_(2,87)_ = 37.48, p < 0.001; males: F_(2,33)_ = 38.20, p < 0.001; center < edge and corner, p < 0.05. Calcium signal was elevated in the center of the arena in both males and females, independent of CIE exposure, in both CeA **(I-J;** 2-way ANOVA, main effect of distance from center, F_(5,175)_ = 27.76, p < 0.001, 0-5 and 6-10 cm from center > rest of bins, p < 0.05) and BNST **(L-M;** main effect of distance from center, F_(5,140)_ = 21.48, p < 0.001, 0-5 cm from center > rest of bins, p < 0.05). Representative calcium traces in CeA **(K)** and BNST **(N)** throughout NSF, with color-coding indicating engagement of discrete behaviors. **O-P:** Data acquired from CeA. **O)** CeA calcium transients during food-approaches reduced across repeated bouts of food-approaches, independent of sex or CIE exposure; 3-way ANOVA on peak z-score, main effect of approach bin only, F_(2,60)_ = 3.85, p = 0.03; bouts 1-5 > bouts 11-16, p < 0.05. **P)** CIE increased CeA calcium signal during the first food-approach in both males and females (2-way ANOVA, main effect of treatment, F_(1,35)_ = 9.20, p = 0.005), and had no effect on calcium signal during sniffing, grooming, and rearing. Females exhibited reduced CeA signal during sniffing behaviors (main effect of sex, F_(1,35)_ = 4.14, p = 0.05). **Q-R:** Data acquired from BNST. **Q)** BNST calcium transients during food-approaches reduced across repeated bouts of food-approaches, independent of sex, with CIE-induced increases in later bouts; 3-way ANOVA on peak z-score, main effect of approach, F_(2,48)_ = 9.93, p = 0.03; bouts 1-5 > bouts 6-10 and 11-15, p < 0.05, interaction effect of approach by treatment, F_(2,48)_ = 4.80, p = 0.01, AIR bout 6-10 > bout 11-15, CIE bout 1-5 < bouts 6-10 and 11-15. **R)** CIE increased BNST calcium signal during the first food-approach in both males and females and had no effect on all other assayed behaviors; 2-way ANOVA, main effect of treatment, F_(1,28)_ = 4.64, p = 0.04.

We observed a sex-specific effect of CIE on latency to feed in females and males. CIE females exhibited an increase in latency to feed relative to alcohol-naïve females **(Fig. 2D)**, whereas CIE males displayed a trend towards a reduction in latency to feed **(Fig. 2C)**. We then employed a machine learning approach to characterize discrete approaches toward the food pellet located in the center zone, as well as ethologically relevant behaviors including sniffing, grooming, and rearing. We found no effect of CIE or sex on the number of food approaches or number of sniffing and grooming bouts prior to food consumption **(Fig. 2E-F)**. CIE females engaged in more rearing prior to eating relative to alcohol-naïve females **(Fig. 2F)**. Time spent in the center zone was lower than time spent in the periphery of the arena in both sexes **(Fig. 2H, G)**, demonstrating thigmotaxis in both sexes; however, time spent in the center and periphery was unaffected by CIE.

We next assessed calcium dynamics in the CeA **(Fig. 2I-K)** and BNST **(Fig. 2L-N)** across the spatial extent of the NSF arena and during discrete behaviors. Calcium signal scaled with spatial proximity to the center zone, independent of sex or CIE treatment, in both the CeA **(Fig. 2J)** and BNST **(Fig. 2M)**. Calcium transients in the CeA and BNST synchronized with the onset of approaches toward the food **(Fig. 2K; 2N)**. To assess whether these food-approach transients were associated with behavioral avoidance, we examined changes in calcium signal across repeated approaches prior to eating. In the CeA, calcium transients decreased across repeated food approaches, independent of sex or treatment **(Fig. 2O)**, suggesting that CeA activity may encode food-related salience or novelty that decreases across repeated approaches, independent of CIE exposure. While the BNST likewise exhibited a decrease in calcium activity across repeated food approaches, CIE exposure heightened BNST calcium activity relative to AIR, independent of sex, demonstrating that CIE prolongs the temporal duration of hyperactivation in the BNST **(Fig. 2Q).**

The onset of food consumption similarly increased calcium activity in the CeA and BNST **(Fig. 2K, 2N)**, and the number of food approaches prior to eating varied across subjects. To avoid confounding eating-associated increases in calcium signal with food-approach transients, we focused our analysis on the first food approach. CeA calcium signal during the first approach was elevated in CIE mice relative to AIR controls, independent of sex **(Fig. 2P)**, demonstrating that CIE promotes hyperactivity in the CeA during the initial food-approach. CIE had no effect on CeA calcium signal across all other ethological behaviors (sniffing, grooming, and rearing), and females exhibited lower calcium signal than males while sniffing food **(Fig. 2P)**. Similar to the CeA, BNST calcium signal was elevated in CIE mice during the first food-approach bout **(Fig. 2R)** independent of sex. We found no effect of CIE or sex on BNST signal during rearing, sniffing, and grooming behaviors **(Fig. 2R)**.

### CIE exposure alters GABAergic synaptic transmission onto BNST neurons in a sex-specific manner

GABA release onto BNST neurons is linked to alcohol self-administration^42,43^ and anxiety.^13,44^ To assess whether our observation of CIE-induced hyperactivity in the BNST of female mice is attributable to impaired inhibitory tone, we used whole-cell patch-clamp electrophysiology to record spontaneous inhibitory postsynaptic currents (sIPSCs) in the BNST of CIE-exposed female mice **(Fig. 3C)**. Given that the CeA is primarily GABAergic,^45^ we selectively assessed GABAergic transmission in this pathway. To selectively stimulate GABAergic inputs from CeA to BNST, we bilaterally injected a Cre-dependent virus encoding the light-gated receptor channelrhodopsin into the CeA of transgenic mice expressing Cre in VGAT-expressing cells **(Fig. 3A)**. 5-7 days following the last CIE exposure, we recorded sIPSCs from BNST neurons. Mean plasma alcohol concentrations per cycle were maintained above binge intoxication levels **(Fig. 3B).** To distinguish BNST neurons within the CeA BNST pathway, we compared cells that were light-responsive (BNST^LR^) to optogenetic stimulation of CeA GABA inputs with cells that were not light-responsive (BNST^non-LR^).

**Fig 3.**
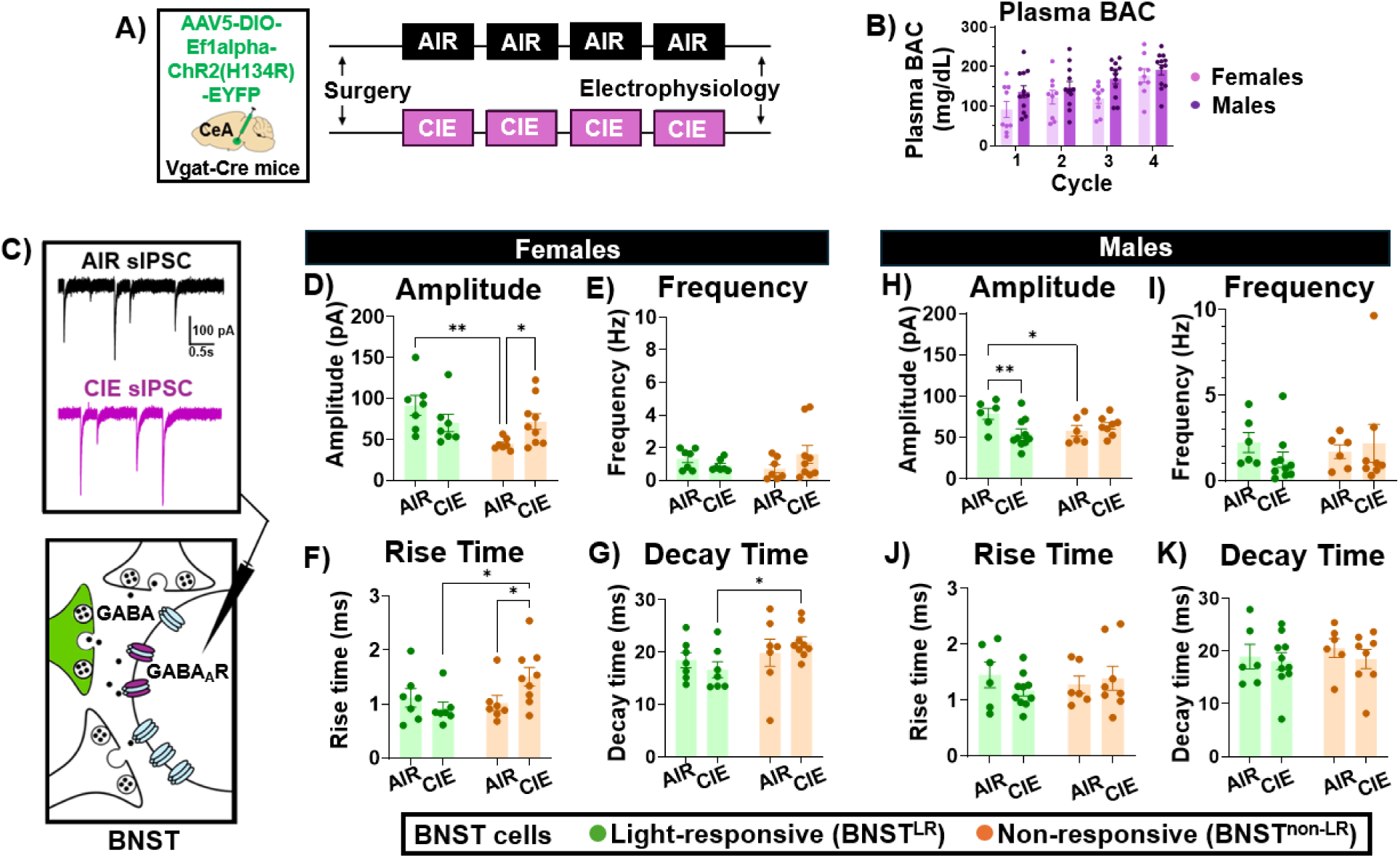
Chronic intermittent ethanol (CIE) exposure alters GABAergic synaptic transmission driven by CeA inputs onto BNST neurons via a sex-specific manner. **A)** Schematic of experimental timeline. Female and male Vgat-Cre mice received bilateral intra-CeA injections of Cre-dependent virus encoding ChR2 prior to CIE exposure. Mice underwent exposure to either vaporized ethanol (CIE; N = 9 female, 10 male) or air (AIR; N = 9 female, 8 male) on a 16hr-ON/8hr-OFF cycle for four weeks. Slice electrophysiological recordings in the BNST were acquired 5-7 days following the last vapor exposure. B) Mean plasma blood alcohol concentrations per cycle were maintained above binge intoxication levels in both females (mean = 127 ± 17.8 mg/dL ; Cycle 1, mean = 91.5 ± 20.2 mg/dL; Cycle 2, mean = 123 ± 17.6 mg/dL; Cycle 3, mean = 119 ± 12.1 mg/dL; Cycle 4, mean = 177 ± 17.0 mg/dL) and males (mean = 161 ± 12.3 mg/dL; Cycle 1, mean = 136 ± 15.2 mg/dL; Cycle 2, mean = 147 ± 15.0 mg/dL; Cycle 3, mean = 170 ± 12.9 mg/dL; Cycle 4, mean = 192 ± 12.7 mg/dL). **C)** Representative sIPSC traces recorded in the BNST of an alcohol-naive control and CIE-exposed mouse (depicted in black and purple, respectively). **D-G:** sIPSC data collected from female mice. N: 7 BNST^L^ cells and 9 BNST^nonLR^ cells from 9 CIE females, 7 BNST^LR^ cells and 7 BNST^nonLR^ cells from 9 AIR females. CIE increased sIPSC amplitude **(D)** and rise time **(F)** in BNST cells that are non-responsive to LED light stimulation; 2-way ANOVA, CIE x light responsivity interaction, F_(1,26)_ = 6.481, p = 0.0172; F_(1,26)_ = 4.709, p = 0.0393. Frequency **(E)** and decay time **(G)** were unaltered by CIE exposure; 2-way ANOVA, all p > 0.05. **H-K:** sIPSC data collected from male mice. N: 10 BNST^LR^ cells and 8 BNST^nonLR^ cells from 10 CIE males, 6 BNST^LR^ cells and 6 BNST^nonLR^ from 8 AIR males. **H)** CIE reduced the amplitude of inhibitory transmission in light-responsive neurons in the BNST; 2-way ANOVA, CIE x light responsivity interaction, F_(1,26)_ = 6.336, p = 0.0183. Frequency **(I)**, rise time **(J)** and decay time **(K)** were unaltered by CIE exposure; 2-way ANOVA, all p > 0.05.

In female mice, CIE increased sIPSC amplitude and rise-time in BNST^non-LR^ neurons with no change in BNST^LR^ neurons, suggesting that CIE selectively enhances GABA signaling from non-CeA originating inputs to the BNST **(Fig. 3D, 3F)**. CIE had no effect on frequency or decay time in females **(Fig. 3E, 3G)**. In males, CIE selectively reduced the amplitude of sIPSCs in BNST^LR^ neurons without altering that of BNST^non-LR^ neurons **(Fig. 3H)**. Frequency, rise-time, and decay-time of BNST sIPSCs were unaltered by CIE in males **(Figs. 3I-3K)** demonstrating that CIE alters GABAergic synaptic transmission onto the BNST of both female and male mice via sex-specific mechanisms. There was no effect of CIE on membrane resistance and capacitance **(Supplementary Table 1)**, suggesting that CIE-induced changes in sIPSC amplitude and kinetics are not attributable to changes in membrane properties.

### CIE exposure speeds the decay of GABA_A_R-mediated synaptic currents driven by CeA inputs to the BNST in female mice

Next, to directly investigate the impact of CIE on synaptic connectivity between CeA and BNST, we used optogenetics to activate and record the CeA→BNST pathway **(Fig. 4C-D)** using slice electrophysiology as described above **(Fig. 3A; 4A)**. Mean plasma blood alcohol concentrations per cycle were maintained above binge intoxication levels **(Fig. 4B).** CIE exposure can disrupt synaptic function via multiple mechanisms, including changes in the subunit composition of postsynaptic GABA_A_Rs and NMDARs, which can alter receptor kinetics.^46,40^ To quantify the decay rate of optically-evoked IPSCs (oIPSCs) in the CeA→BNST pathway, we fitted averaged oIPSC traces **(Fig. 4D)** to a single exponential decay. CIE reduced the decay rate of oIPSCs in this pathway in female but not male mice **(Fig. 4E)**. These findings demonstrate that CIE promotes hyperactivation in the BNST of female mice by speeding the decay of GABA_A_R-mediated synaptic currents in the CeA BNST pathway.

**Fig 4.**
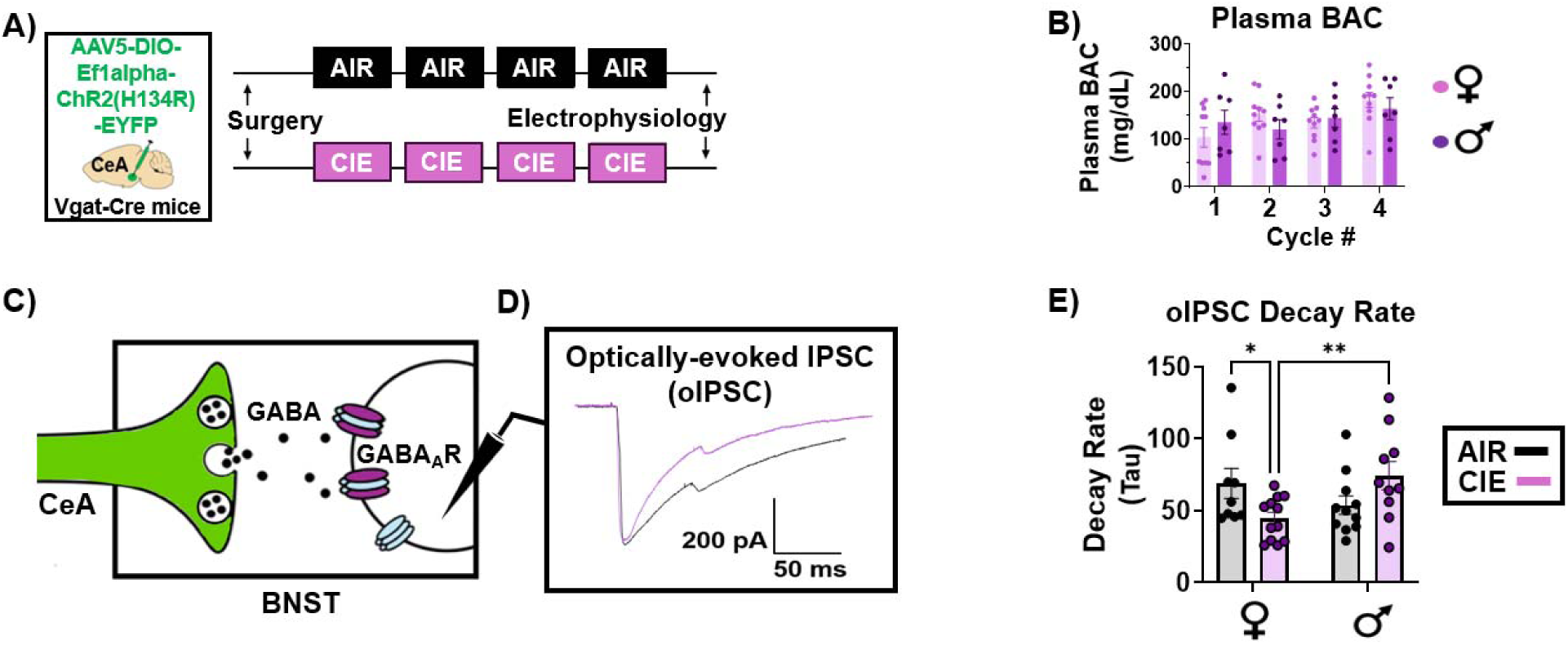
CIE exposure speeds the decay of GABA_A_-R mediated synaptic currents driven by CeA inputs to the BNST of female mice. **A)** Schematic of experimental timeline. Female and male Vgat-Cre mice received bilateral intra-CeA injections of Cre-dependent virus encoding ChR2 prior to CIE exposure. Mice underwent exposure to either vaporized ethanol (CIE; N = 10 female, 8 male) or air (AIR; N= 6 female, 8 male) on a 16hr-ON/8hr-OFF cycle for four weeks. Slice electrophysiological recordings in the BNST were acquired 5-7 days following the last vapor exposure. **B)** Mean plasma blood alcohol concentrations per cycle were maintained above binge intoxication levels in both females (mean = 144 ± 16.4 mg/dL; Cycle 1, mean = 105 ± 20.3 mg/dL; Cycle 2, mean = 152 ± 14.2 mg/dL; Cycle 3, mean = 135 ± 11.2 mg/dL; Cycle 4, mean = 183 ± 16.1 mg/dL) and males (mean = 141.8 ± 9.08 mg/dL; Cycle 1, mean = 136 ± 25.4 mg/dL; Cycle 2, mean = 121.2 ± 20.3 mg/dL; Cycle 3, mean = 145 ± 19.2 mg/dL; Cycle 4, mean = 165 ± 22.9 mg/dL). **C)** Schematic of experimental design for obtaining optically-evoked IPSC recordings in BNST neurons during optogenetic stimulation of CeA GABAergic inputs to the BNST during baseline conditions. **D)** Representative oIPSC traces from an alcohol-naïve female mouse (depicted in black) versus a CIE-exposed female mouse (depicted in purple). **E)** CIE reduced the decay rate of GABA_A_-R mediated synaptic currents driven by CeA inputs to the BNST of female mice, but had no effect in male mice; 2-way ANOVA, CIE x sex interaction, F_(1,38)_ = 8.420, p = 0.0061. N: 12 cells from 10 CIE females and 9 cells from 6 AIR females, 10 cells from 8 CIE males and 11 cells from 8 AIR males.

### CIE exposure increased KOR suppression of the CeA BNST pathway in a sex-specific manner

KORs in the CeA and BNST have been heavily implicated in alcohol induced anxiety^29–32,31^ and presynaptic KORs located in the axonal terminals of the CeA suppress GABA release from CeA inputs to the BNST in alcohol-naïve males **(Fig. 5C)**.^33^ Thus, we examined the KOR agonist U69593 (1µM) on oIPSCs in the CeA→BNST pathway following CIE exposure **(Fig. 5A, 5C)**. Mean plasma blood alcohol concentrations per cycle were maintained above binge intoxication levels **(Fig. 5B).** In both male and female control mice, we found that KOR agonist reduced peak amplitude of oIPSC **(Fig. 5D, 5F)**. Given that CIE exposure upregulates KOR signaling in both the CeA and BNST,^29,31,47^ we hypothesized that CIE enhances KOR signaling in the CeA→BNST pathway, leading to reduced GABA release during alcohol withdrawal. In females, CIE exposure had no effect on the degree of KOR-induced suppression of oIPSC amplitude, contrary to our hypothesis that CIE upregulates KOR signaling in this pathway **(Fig. 5E)**. However, in males we observed that CIE exposure increased the extent of KOR-induced suppression of oIPSC amplitude **(Fig. 5G)**.

**Fig 5.**
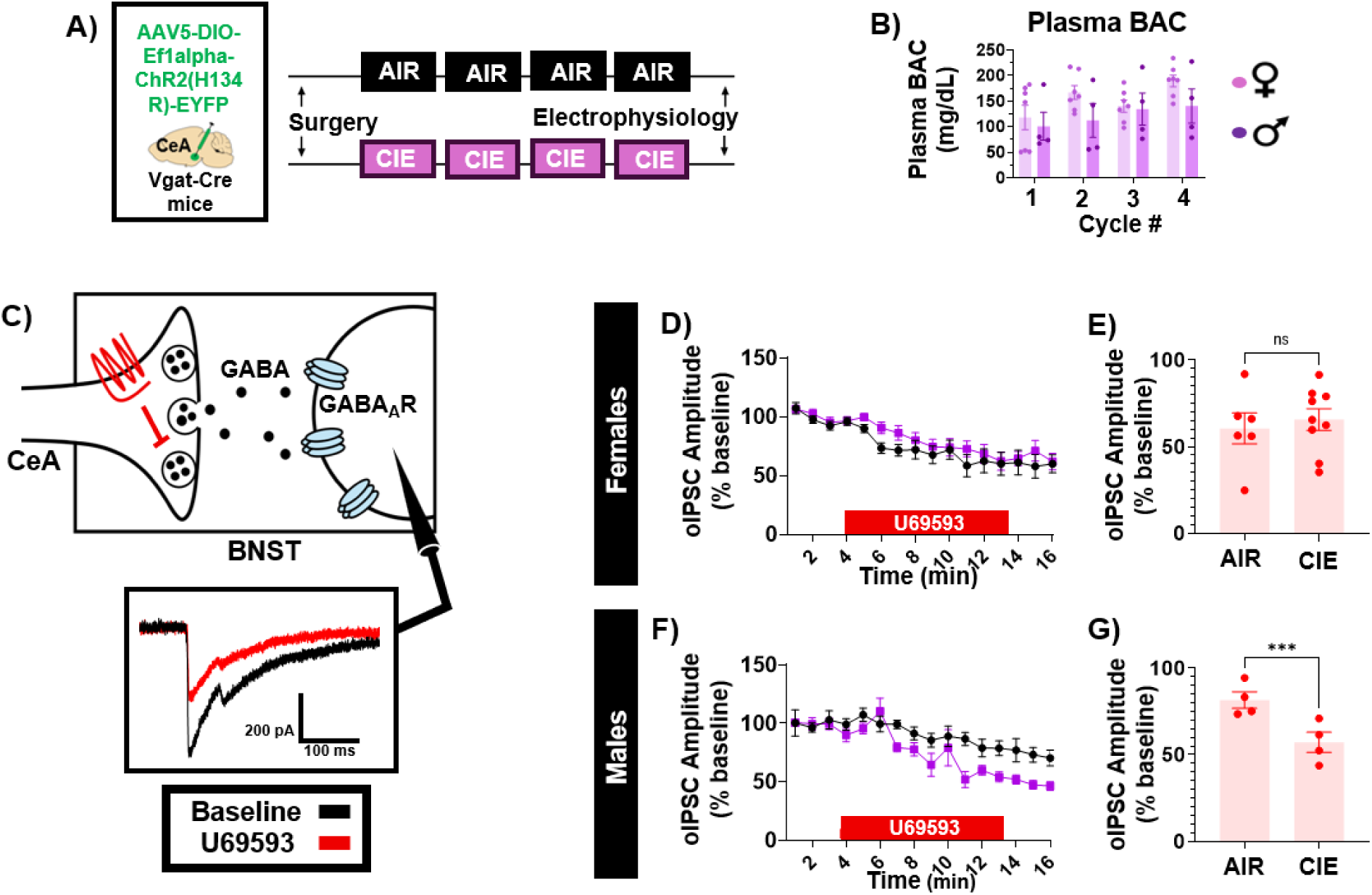
CIE exposure increased KOR suppression of the CeA BNST pathway in male but not female mice. **A)** Schematic of experimental timeline. Female and male Vgat-Cre mice received bilateral intra-CeA injections of Cre-dependent virus encoding ChR2 prior to CIE exposure. Mice underwent exposure to either vaporized ethanol (CIE; N = 7 female, 4 male) or air (AIR; N = 5 female, 3 male) on a 16hr-ON/8hr-OFF cycle for four weeks. Slice electrophysiological recordings in the BNST were acquired 5-7 days following the last vapor exposure. **B)** Mean plasma blood alcohol concentrations were maintained above binge intoxication levels in both females (mean = 154 ± 15.6 mg/dL; Cycle 1, mean = 118 ± 23.8 mg/dL; Cycle 2, mean = 167 ± 13.7 mg/dL; Cycle 3, mean = 140 ± 11.8 mg/dL; Cycle 4, mean = 190 ± 11.3 mg/dL) and males (mean = 122 ± 9.29 mg/dL; Cycle 1, mean = 101 ± 27.3 mg/dL; Cycle 2, mean = 112 ± 33.1 mg/dL; Cycle 3, mean = 134 ± 31.5 mg/dL; Cycle 4, mean = 140 ± 33.3 mg/dL). **C)** Representative oIPSC traces recorded from BNST neurons during optogenetic stimulation of CeA inputs to the BNST during baseline conditions and KOR activation as induced by infusion of the KOR agonist U69593 (1µM). **D-E:** oIPSC data collected from female mice. N: 9 cells from 7 CIE females, 6 cells from 5 AIR females. **D)** U69593 suppressed oIPSC peak-amplitude in both AIR controls and CIE-exposed females. **E)** CIE had no effect on KOR suppression; 2-tailed unpaired t-test, t_(9.640)_ = 0.4750, p = 0.6454. **F-G:** oIPSC data collected from male mice. N: 4 cells from 4 CIE males, 4 cells from 3 AIR males. **F)** U69593 suppressed CeA BNST inhibitory transmission in AIR controls and CIE-exposed males. **G)** CIE exposure increased the degree of KOR suppression; 2-tailed unpaired t-test, t_(5.760)_ = 3.230, p < 0.0190.

## Discussion

Here, we demonstrate that CIE promotes increased anxiety-like behavior in female mice and BNST hyperactivity during approach behavior in anxiogenic contexts. Our circuit-specific electrophysiological studies elucidated a potential mechanism for this hyperactivity, identifying a female-specific postsynaptic disruption in the CeA→BNST pathway during alcohol abstinence. We also observed a frequency-specific increase in CeA-BNST coherence in CIE-exposed females during open arm entries in the EPM, suggesting enhanced functional coupling between these structures during anxiogenic tasks. Altogether, these observations highlight discrete, sex-specific cellular adaptations in the CeA→BNST circuit that may contribute to heightened BNST activity during alcohol withdrawal and represent potential therapeutic targets for mitigating alcohol-induced anxiety in women with AUD.

### Female mice exhibit increased BNST activity during anxiety-like states following alcohol dependence

We found that CIE increased avoidance in females as measured in both the NSF and EPM. This anxiety-like behavioral phenotype in CIE females was accompanied by calcium activity changes in BNST suggestive of increased excitability. Using Hidden Markov Models to identify distinct behavioral states in the EPM, we found that CIE promotes hyperactivity in the BNST during surveillance behaviors (State 2), walking (State 3), and when not occupying a distinct behavioral state. BNST calcium activity was increased during deceleration and reduced during acceleration, suggesting neuronal activity is inversely related to locomotor activity independent of CIE. CIE potentiated the increase in BNST activity during deceleration and attenuated the suppression in BNST activity during acceleration, suggesting that BNST hyperactivity may suppress movement initiation. Altogether, our observations that CIE reduces locomotion and increases freezing behavior are consistent with increased behavioral inhibition and support a role for BNST hyperactivity during anxiety-like states. To our knowledge, our findings provide the first direct *in vivo* evidence that chronic alcohol increases BNST activity during anxiety-related behaviors. Interestingly, we observed sex differences in the effects of CIE on latency to feed in the NSF assay. Whereas CIE females displayed an anxiogenic phenotype, in keeping with previous findings by Centanni et al,^22^ CIE males showed a trend toward reduced latency, contrasting other work.^48^ This may be explained by differences in the abstinence timepoint, duration of fasting, and type of food presented in the test.

### CIE reduced GABA tone in the CeA→BNST pathway in a female-specific manner

Consistent with the female-specific BNST hyperactivity observed following CIE, CIE sped up the decay of GABA_A_-R mediated synaptic currents in the CeA→BNST pathway in females but not males. This change in decay kinetics following CIE is in keeping with previous reports in the hippocampus.^39^ The decay kinetics of sIPSCs in BNST neurons were unaltered by CIE, suggesting input-specific changes rather than a global adaptation within the BNST. Relevant to AUD, our observation that CIE reduces GABA tone in this pathway in a female-specific manner is in line with clinical evidence demonstrating sex-specific alterations in BNST circuitry in AUD, including stronger structural BNST network connectivity in women than in men.^14^

To address gaps in knowledge regarding how CIE affects synaptic physiology in the BNST of female mice, we recorded spontaneous IPSCs in the BNST following CIE. In accordance with previous findings in males, CIE had no effect on inhibitory drive onto BNST neurons, independent of sex. When analyzing in an optogenetic projection-specific manner, however, we observed that CIE potentiated GABA transmission onto BNST neurons that ostensibly lack CeA inputs (BNST^non-LR^), as measured by increased amplitude and rise-time. Although a stronger inhibitory drive onto BNST neurons seems counterintuitive given heightened BNST activity during alcohol-induced anxiety, previous work by Marcinkiewcz et al demonstrated that a subpopulation of midbrain-projecting cells in the BNST exhibit reduced activity during anxiety-like behaviors, suggesting the effects of CIE may be subpopulation specific.^49^ In male mice, CIE reduced the sIPSC amplitude of BNST neurons that ostensibly receive CeA inputs, suggesting a male-specific postsynaptic adaptation to the CeA→BNST pathway. This suggests that CIE promotes BNST hyperactivity in males by selectively disrupting GABA signaling at the CeA→BNST synapse.

### KORs suppress GABA release from CeA inputs to BNST in female mice

Our observations provide the first evidence that the CeA→BNST pathway is sensitive to KOR signaling in females, potentially driving disinhibition of BNST neurons in female mice. This disinhibition may enable synchronous CeA-BNST activation by common upstream inputs. Relevant to AUD, our observation of stronger frequency-specific coherence between CeA and BNST during anxiety-like states following CIE aligns with human neuroimaging data demonstrating that stronger CeA-BNST functional connectivity positively correlates with higher trait anxiety measures and amygdala-BNST connectivity is increased during alcohol abstinence in individuals with AUD.^11,15^ To our knowledge, no clinical or preclinical studies to date have examined CeA-BNST functional connectivity following alcohol exposure, nor the impact of sex as a statistical variable in anxiety studies of CeA-BNST functional connectivity. Our data suggests that this may be a useful biomarker for exploring anxiety-states.

### Alcohol dependence enhanced the modulatory gain of KORs in the CeA→BNST pathway via a sex-specific manner

In accordance with our hypothesis that CIE likely exacerbates the extent of KOR suppression in the CeA→BNST pathway, we found that CIE potentiated the extent of this KOR suppression in male mice. To our surprise, we found no effect of CIE on this KOR suppression in females. Our findings demonstrate that CIE increased the capacity of KORs to suppress GABA transmission in this pathway, and the male-specificity of this effect aligns with previous work by Bloodgood at al demonstrating that genetic deletion of CeA KORs following a history of alcohol consumption reduces alcohol self-administration in males but not females.^50^ One key limitation of our findings is that we did not perform EPM in male mice, precluding a more robust sex comparison. Our findings further our understanding of the effects of alcohol on the female extended amygdala and could inform the future development of pharmacotherapeutic and diagnostic strategies to mitigate alcohol-induced anxiety in women with AUD. Future studies exploring synapse dependent changes in GABA_A_-R function in the BNST may identify novel pharmacological approaches to explore alcohol driven negative affect.

## Author contributions

**Sarah Chong, Alison V. Roland, Jobe L. Ritchie**: Conceptualization, Writing – original draft, Investigation, Formal Analysis. **Lisa R. Taxier, Sloan Markowitz, Lili S. Kooyman, Maya Eberle-Redding, Chloe Gallagher, Madison R. Campeau:** Writing – review & editing, Investigation. **Thomas L. Kash**: Supervision, Conceptualization, Writing – review & editing, Funding acquisition.

## Supplementary Information

### Drug Information

Recording electrodes were filled with (in mmol/L) 117 Cs-Gluc, 20 HEPES, 0.4 EGTA, 5 TEA, 2 MgCl2, 4 Na2ATP, 0.4 Na2GTP. Lidocaine N-ethyl bromide (1mg/mL) was included in the intracellular recording solution to prevent postsynaptic sodium spikes. Both spontaneous inhibitory postsynaptic currents (sIPSCs) and optically-evoked inhibitory postsynaptic currents were pharmacologically isolated by adding 3 mmol/L kynurenic acid to block alpha-amino-3-hydroxy-5-metnyl-4-isocsazole-propionic acid (AMPA) and N-methyl-D-aspartate (NMDA) receptor-dependent postsynaptic currents. (+)-(5α,7α,8β)-N-Methyl-N-[7-(1-pyrrolidinyl)-1-oxaspiro[4.5]dec-8-yl] benzeneacetamide (U69593, 1 µM), a synthetic agonist highly selective for the kappa opioid receptor (La Regina et al, 1988), was purchased from Sigma-Aldrich (St. Louis, MO) and dissolved in 0.1 M HCl.

### Supplementary Table

**Supplementary Table 1.**
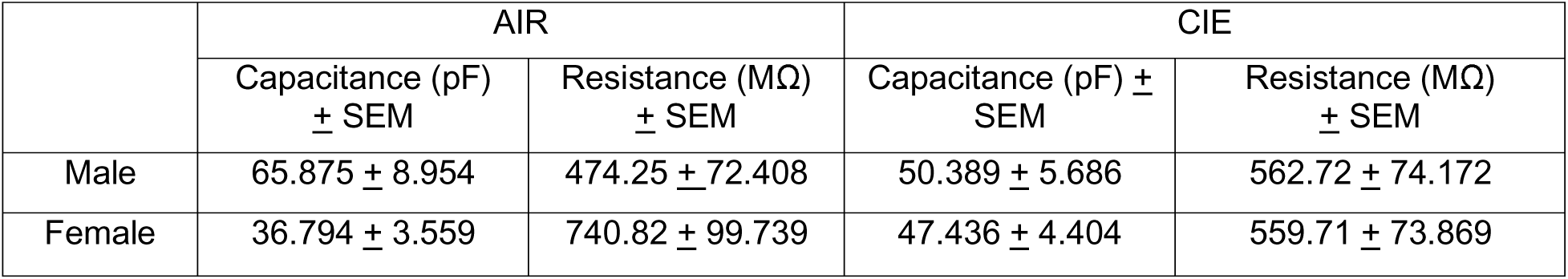
Cell Properties in sIPSC Experiments.

|  | AIR |  | CIE |  |
|---|---|---|---|---|
|  | Capacitance (pF) ± SEM | Resistance (MΩ) ± SEM | Capacitance (pF) ± SEM | Resistance (MΩ) ± SEM |
| Male | 65.875 ± 8.954 | 474.25 ± 72.408 | 50.389 ± 5.686 | 562.72 ± 74.172 |
| Female | 36.794 ± 3.559 | 740.82 ± 99.739 | 47.436 ± 4.404 | 559.71 ± 73.869 |

